# Raspberry pi: Assessments of emerging organic chemicals by the predictive *in silico* methods

**DOI:** 10.1101/2021.01.15.426465

**Authors:** Charli Deepak Arulanandam, R Prathiviraj, Govindarajan Rasiravathanahalli Kaveriyappan

## Abstract

Phthalic acid esters (PAEs) and bisphenols are used as plasticizers worldwide. During plastic production, use, deposition, and recycling these compounds contaminate the environment and affect environmental health. In this study, we investigated the toxicity of plasticizers by using *in silico* tools. None of the test compounds were found to be hERG blockers in multiclass predictions as evaluated by the Pred-hERG 4.1 tool. Among all tested compounds in Pred-Skin 2.0, only BBP, BCP, DBP, diethyl phthalate (DEP), DMP, DNHP, DNPP, DPP, DTDP, DUP, and ODP were non-skin sensitizers. Our results demonstrate that *in silico* tools provide a reliable, fast, and economic way to explore the toxicological effects of EOCs.

## Introduction

Plastic products are used for different purposes in everyday life. These provide several prospects for the users, medical care, and industrial progressions (Thompson *et al*., 2009). At the same time possibility of the plastic contaminants in food products from the food-packaging materials lead to human health risks (Claudio, 2012; Liao and Kannan, 2013). Plastic waste in landfills, as well as microplastic discharge into the ground, coastal, and ocean waters. Plasticizers that stabilize or soften plastics are hydrophilic and soluble at the same time, they may contaminate aquatic environments. If they are volatile, they pollute the atmosphere and may dissolve in water vapor that precipitates eventually or affects the human respiratory organ. Plasticizers are emerging organic chemicals (EOCs) that were detected in the environment (Vandenberg *et al*., 2007). These widely distributed environmental contaminants with toxic effects on animals and humans, (Sarath *et al*., 2016) were even measured in marine aerosol samples over the East China Sea among other organic aerosols (OA), due to the influence of terrestrial emissions on oceanic aerosols from fossil fuel combustion (Kang *et al*., 2017). Among other effects are EOCs causing endocrine disruption in animals and humans. If being lipophilic, Phthalates can easily be absorbed and diffused in lipids, causing physiological dysfunctions and ontogenetic bioaccumulation with age and biomagnification within food webs (Benjamin *et al*., 2015). Bisphenols are phenol derivatives (Fiege *et al*., 2000). They are used for the synthesis of epoxy resins, polycarbonates, and thermal paper. The annual production of bisphenol A (BPA) alone exceeds 3.8 million tons (Hoekstra and Simoneau, 2013). It commonly appears in several products of everyday use including water-pipes, electronic equipment, paper, and toys (Flint *et al*., 2012). There is increasing evidence that BPA is harmful to human health (Rochester, 2013). Looking at animal models, rodent exposure to BPA affects particularly the reproductive functioning of males. Long-lasting effects upon chronic BPA exposure also affect metabolic processes in males and females of rodent animal models (Richter *et al*., 2007). BPA is the most important anthropogenic EDC. It affects the functions of several hormones. Since BPA can leach into food and water supplies, it provides a particular threat to drinking water and food safety (Niu *et al*., 2012). This resulted in considerable research into exposure-associated health risks in humans. The removal of BPA from several plastic products that followed consumer alerts caused an increasing development and use of bisphenol analogues (Clark, 2000). As for governmental regulations, chemicals should be replaced by less toxic or non-toxic chemicals to avoid adverse effects on consumers. However, several chemical replacements remain untested before being marketed and might be toxic or even more harmful than the original ones. Experimental screening of chemical compounds for biological activity by *in situ* and *in vitro* experiments is an expensive practice in terms of investing materials and expert efforts in space and time. In this situation could *in silico* predictive models provide inexpensive alternatives (Seal *et al*., 2012). Computational approaches can also provide priority criteria for subsequent *in vitro* testing of toxicological evaluations. This will reduce and replace experiment costs and animal testing (McRobb *et al*., 2014). We want to emphasize here that chemically transformed compounds of EOCs such as BPA are only rarely considered in environmental risk assessments as yet (Gao *et al*., 2015). Application of *in silico* tools would reduce animal testing and increase the effectiveness of risk assessments (Worth *et al*., 2011). In this study, we used various predictive tools and a web servers for the evaluations of EOCs. In this assessment, we evaluated hepatotoxicity, carcinogenicity, cardiac toxicity, and skin sensitization. We hypothesize that *in silico* methods provide useful tools for the predictions of different toxicological endpoints. Here, we particularly screened for possible toxicities of plasticizers with help of Pred-hERG 4.1, and Pred-Skin 2.0.

## Materials and Methods

### Raspberry pi System configuration

Raspberry Pi 3 Model B+ installed with raspbian operating system (OS) and this machine utilized for this *in silico* approach. The speed and performance of the Raspberry Pi 4 is a step up from earlier models. This hardware has been selected for highly reliable technology in the compact machine but more energy-efficient and much more cost-effective machine. The system configuration of the used machine is SoC: Broadcom BCM2837B0 quad-core A53 (ARMv8) 64-bit @ 1.4GHz, GPU: Broadcom Videocore-IV, RAM: 1GB LPDDR2 SDRAM, Networking: Gigabit Ethernet, 2.4GHz and 5GHz 802.11b/g/n/ac Wi-Fi, Bluetooth: Bluetooth 4.2, Bluetooth Low Energy (BLE), Storage: Micro-SD, GPIO: 40-pin GPIO header, populated, Ports: HDMI, 3.5mm analogue audio-video jack, 4x USB 2.0, Ethernet, Camera Serial Interface (CSI), Display Serial Interface (DSI), Dimensions: 82mm x 56mm x 19.5mm, 50g for more detail regarding this machine visit to the https://www.raspberrypi.org.

### Data collection and retrieval

Plasticizers were retrieved from ChemSpider and PubChem are summarized with abbreviations in STable 1 and 2). The molecular two-dimensional structures of plasticizers were drawn using Marvin 17.21.0, ChemAxon (Fig. 1), based on available molecular data from ChemSpider. This Java-based chemical drawing tool allowed to edit molecules in different file formats available from https://www.chemaxon.com/products/marvin (see also Evans and Moore, 2011).

**Table 1.**
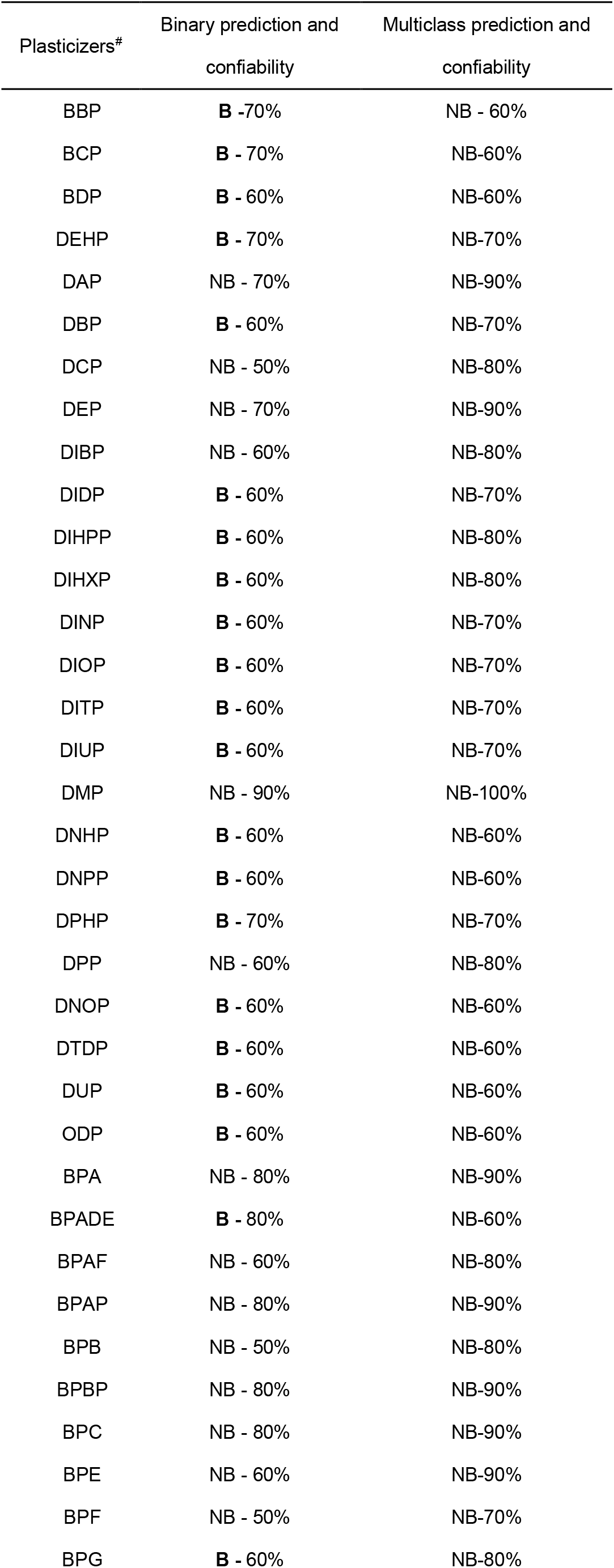
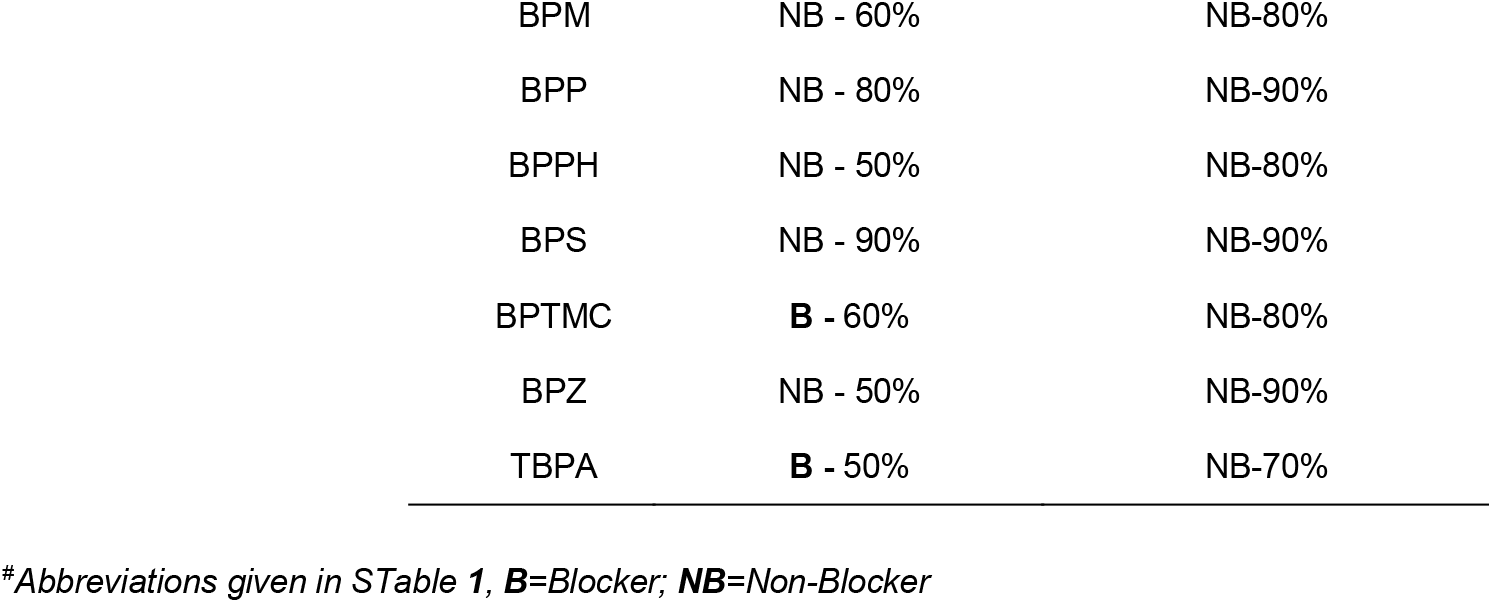
hERG blocking plasticizers predicted by Pred-hERG 4.1 tool.

**Table 2.** Skin sensitizing property of plasticizers.

| **Table 2.** Skin sensitizing property of plasticizers. |  |  |  |
| --- | --- | --- | --- |
| Plasticizers<sup>#</sup> | LLNA model |  | Human skin model |
|  | Binary prediction | Multiclass prediction | Binary prediction |
| BBP | NS | NS | NS |
| BCP | NS | NS | NS |
| BDP | NS | W/M | NS |
| DEHP | NS | NS | NS |
| DAP | S | NS | NS |
| DBP | NS | NS | NS |
| DCP | S | NS | NS |
| DEP | NS | NS | NS |
| DIBP | S | NS | NS |
| DIDP | S | W/M | NS |
| DIHPP | S | NS | NS |
| DIHXP | S | NS | NS |
| DINP | S | W/M | NS |
| DIOP | S | W/M | NS |
| DITP | S | W/M | NS |
| DIUP | S | W/M | NS |
| DMP | NS | NS | NS |
| DNHP | NS | NS | NS |
| DNPP | NS | NS | NS |
| DPHP | S | NS | NS |
| DPP | NS | NS | NS |
| DNOP | S | NS | NS |
| DTDP | NS | NS | NS |
| DUP | NS | NS | NS |
| ODP | NS | NS | NS |
| BPA | S | W/M | NS |
| BPADE | S | W/M | S |
| BPAF | S | W/M | S |
| BPAP | S | NS | NS |
| BPB | S | NS | S |
| BPBP | S | NS | S |
| BPC | S | W/M | S |
| BPE | S | NS | S |
| BPF | S | S/E | S |
| BPG | S | W/M | S |
| BPM | S | NS | NS |
| BPP | S | W/M | NS |
| BPPH | S | W/M | S |
| BPS | S | NS | S |
| BPTMC | S | NS | NS |
| BPZ | S | S/E | S |
| TBPA | S | W/M | S |

<sup>#</sup>Abbreviations given in STable 1, **NS**=Non-Sensitizer, **S**=Sensitizer; **W/M**=Weak/Moderate; **S/E**=Strong/Extreme

### SMILES based input format for compounds

Molecular structures are represented as SMILES (simplified molecular-input line-entry system) which were needed as 2D-input data for the SMILES based similarity functions are computationally more efficient in an *in silico* approach (Frenzel *et al*., 2017). SMILES of the test compounds are shown in **STable 3**.

### Cardiac toxicity prediction

Bisphenols may contribute to cardiometabolic risks as well and childhood exposure to bisphenols affecting child growth (Philips *et al*., 2017). The inhibition of human ERG (an ether-a-go-go-related gene synthesizing an enzyme that blocks K+ channels). This inhibition leads to heart arrhythmia that can cause death. Even non-cardiovascular drugs were withdrawn from the market upon this discovery. An hERG safety screening became then mandatory following a procedure required by the food and drug administration (FDA) in the USA (Braga et al. 2014). The Pred-hERG 4.1 web server predicts potential hERG blockers and non-blockers from different chemicals (Braga et al. 2015) and was applied to predict the cardiac toxicity of plasticizers. This web server tool is available at http://labmol.com.br/predherg/.

### Skin sensitization toxicity prediction

Skin sensitization represents an immunological disorder. This webserver was developed based on binary QSAR models predicting skin sensitization according to a local lymph node assay (LLNA) derived from experimental data from rodents (=murine) and human. Braga and co-workers (2017) included a multiclass skin sensitization potency model based on LLNA data. When a user evaluates a compound in the web app, the output is (i) binary predictions of human and murine skin sensitization potential; (ii) a multiclass prediction of murine skin sensitization; and (iii) probability maps illustrating the predicted contribution of chemical fragments. The Pred-Skin web app version 2.0 is available from the web, iOS at the LabMol portal, in the Apple Store, and on Google Play, respectively (Braga *et al*., 2017). Such a tool is used to predict the skin sensitization of plasticizers and is accessed at http://labmol.com.br/predskin/.

## Results and Discussion

From the Pred-hERG 4.1 cardiac toxicity assessed by using the SMILES of plasticizers. This web server prediction shows as none of the tested plasticizers blocking the hERG (see Table 1). However, in vitro studies demonstrated that acute BPA exposure caused heart arrhythmia in female rodents and BPA showed cardiovascular (CV) activity and toxicity (Gao and Wang, 2014). Pred-Skin 2.0 predicted that all tested compounds provided a risk of skin sensitization except BBP, BCP, DEHP, DBP, DMP, DNHP, DNPP, DPP, DTDP, DUP, ODP according to this tool (see Table 2).

## Conclusions

In silico predictive models provide fast and economic screening tools for different other compound properties. Evaluation of plasticizers as EOCs to reduce the amount of costly in vivo and in vitro toxicological testing and also to provide early alerts for environmental and health risk assessment. Here utilized web server freely accessible, fast, predictive, economic, and reliable.

## Acknowledgment

Marvin was used for drawing, displaying, and to obtain SMILES of chemical structures (Marvin17.21.0; ChemAxon (https://www.chemaxon.com) for academic research licence.

## Funding Statement

This work was supported by an International Student KMU-Kaohsiung Medical University fellowship to Arulanandam Charli Deepak.

## Conflict of Interest

All authors agree in having no competing interests concerning this contribution.

## Notes

https://chemdata.org/

### Competing Interest Statement

The authors have declared no competing interest.

