## Supporting Information for "Raspberry pi: Assessments of emerging organic chemicals by the predictive *in silico* methods"

**STable 1**. Abbreviations used

| | |
|---|---|
| **ADMET** | Absorption, Distribution, Metabolism, Excretion, Toxicity |
| **BBP** | benzyl butyl phthalate |
| **BPA** | bisphenol A |
| **BPAF** | bisphenol AF |
| **BPB** | p,p'-sec-Butylidenediphenol /bisphenol B |
| **BPC** | dihydroxymethoxychlor olefin/bisphenol C |
| **BPE** | 4,4'-Ethylidenediphenol/bisphenol E |
| **BPF** | bisphenol F |
| **BPS** | bisphenol S |
| **BPZ** | 4,4'-cyclohexane-1,1-diyldiphenol/bisphenol Z |
| **CarcinoPred-EL** | carcinogenicity prediction using ensemble learning methods |
| **CSIDs** | chemspider identifier |
| **DBP** | dibutyl phthalate |
| **DEHP** | di-(2-ethylhexyl)phthalate |
| **DEP** | diethyl phthalate |
| **DMP** | dimethyl phthalate |
| **DNOP** | dioctyl phthalate |
| **DoTS** | docking interface for Target Systems |
| **EDCs** | endocrine-disrupting chemicals |
| **EOCs** | emerging organic contaminants |
| **hERG** | human ether-a-go-go-related gene |
| **LAZAR** | lazy structure-activity relationships |
| **LD50** | lethal dose, 50% of population |
| **LLNA** | murine local lymph node assay |
| **OA** | organic aerosols |
| **PAEs** | Phthalic acid esters/ Phthalates |
| **QSAR** | quantitative structure-activity relationship |
| **REACH** | Registration, Evaluation, Authorisation and restriction of Chemicals |
| **RF** | random forest |
| **SMILES** | simplified molecular input line entry system |
| **SVM** | support vector machine |
| **VEGA** | virtual models for property evaluation of chemicals within a global architecture |
| **XGBoost** | extreme gradient boosting |

**STable 2.** Plasticizer compound data

| Plasticizers | Abbreviations | Molecular Formula | Chemical ID |
|---|---|---|---|
| benzyl butyl phthalate | BBP | $C_{19}H_{20}O_4$ | CSID 2257 |
| butyl cyclohexyl phthalate | BCP | $C_{18}H_{24}O_4$ | CSID 6521 |
| butyl decyl phthalate | BDP | $C_{22}H_{34}O_4$ | CSID 6697 |
| di-(2-ethylhexyl)phthalate | DEHP | $C_{24}H_{38}O_4$ | CSID 5414319 |
| diallyl phthalate | DAP | $C_{14}H_{14}O_4$ | CSID 8242 |
| dibutyl phthalate | DBP | $C_{16}H_{22}O_4$ | CSID 13837319 |
| dicyclohexyl phthalate | DCP | $C_{20}H_{26}O_4$ | CSID 6519 |
| diethyl phthalate | DEP | $C_{12}H_{14}O_4$ | CSID 13837303 |
| diisobutyl phthalate | DIBP | $C_{16}H_{22}O_4$ | CSID 6524 |
| diisodecyl phthalate | DIDP | $C_{28}H_{46}O_4$ | CSID 30996 |
| diisoheptyl phthalate | DIHPP | $C_{22}H_{34}O_4$ | CSID 513939 |
| diisohexyl phthalate | DIHXP | $C_{20}H_{30}O_4$ | CSID 8623 |
| diisononyl phthalate | DINP | $C_{26}H_{42}O_4$ | CSID 513622 |
| diisooctyl phthalate | DIOP | $C_{24}H_{38}O_4$ | CSID 31280 |
| diisotridecyl phthalate | DITP | $C_{34}H_{58}O_4$ | CSID 141908 |
| diisoundecyl phthalate | DIUP | $C_{30}H_{50}O_4$ | CSID 153056 |
| dimethyl phthalate | DMP | $C_{10}H_{10}O_4$ | CSID 13837329 |
| di-n-hexyl phthalate | DNHP | $C_{20}H_{30}O_4$ | CSID 6528 |
| di-n-pentyl phthalate | DNPP | $C_{18}H_{26}O_4$ | CSID 8243 |
| bis-(2-propylheptyl)phthalate | DPHP | C28H46O4 | CID 92344 |
| di-n-propyl phthalate | DPP | $C_{14}H_{18}O_4$ | CSID 8241 |
| dioctyl phthalate | DNOP | $C_{24}H_{38}O_4$ | CSID 8043 |
| ditridecyl phthalate | DTDP | $C_{34}H_{58}O_4$ | CSID 8076 |
| diundecyl phthalate | DUP | $C_{30}H_{50}O_4$ | CSID 18193 |
| n-Octyl n-decyl phthalate | ODP | $C_{26}H_{42}O_4$ | CSID 8077 |
| bisphenol A | BPA | $C_{15}H_{16}O_2$ | CSID 6371 |
| bisphenol A Diglycidyl Ether | BPADE | $C_{21}H_{24}O_4$ | CSID 2199 |
| bisphenol AF | BPAF | $C_{15}H_{10}F_6O_2$ | CSID 66498 |
| bisphenol AP | BPAP | $C_{20}H_{18}O_2$ | CSID 541979 |
| bisphenol B | BPB | $C_{16}H_{18}O_2$ | CSID 59553 |
| bisphenol BP | BPBP | $C_{25}H_{20}O_2$ | PSID 329748555 |
| bisphenol C | BPC | $C_{14}H_{10}Cl_2O_2$ | CSID 76387 |
| bisphenol E | BPE | $C_{14}H_{14}O_2$ | CSID 528599 |
| bisphenol F | BPF | $C_{13}H_{12}O_2$ | CSID 11614 |
| bisphenol G | BPG | C21H28O2 | CID 228537 |
| bisphenol M | BPM | $C_{24}H_{26}O_2$ | CSID 2540817 |

| | | | |
|---|---|---|---|
| bisphenol P | BPP | $C_{24}H_{26}O_2$ | CSID 547401 |
| bisphenol PH | BPPH | $C_{27}H_{24}O_2$ | PSID 329758221 |
| bisphenol S | BPS | $C_{12}H_{10}O_4S$ | CSID 6374 |
| bisphenol TMC | BPTMC | $C_{21}H_{26}O_2$ | CID 4134035 |
| bisphenol Z | BPZ | $C_{18}H_{20}O_2$ | CSID 202599 |
| tetrachlorobisphenol A | TBPA | C15H12Cl4O2 | CID 6619 |

*CSID - Chemspider Identifier number, CID- PubChem Compound ID, SID-PubChem Substance ID.*

**STable 3**. SMILES of Plasticizers.

| Plasticizers | SMILES |
|---|---|
| BBP | CCCCOC(=O)C1=CC=CC=C1C(=O)OCC1=CC=CC=C1 |
| BCP | CCCCOC(=O)C1=CC=CC=C1C(=O)OC1CCCCC1 |
| BDP | CCCCCCCCCCOC(=O)C1=CC=CC=C1C(=O)OCCCC |
| DEHP | [H][C@](CC)(CCCC)COC(=O)C1=CC=CC=C1C(=O)OC[C@]([H])(CC)CCCC |
| DAP | C=CCOC(=O)C1=CC=CC=C1C(=O)OCC=C |
| DBP | CCCCOC(=O)C1=CC=CC=C1C(=O)OCCCC |
| DCP | O=C(OC1CCCCC1)C1=CC=CC=C1C(=O)OC1CCCCC1 |
| DEP | CCOC(=O)C1=CC=CC=C1C(=O)OCC |
| DIBP | CC(C)COC(=O)C1=CC=CC=C1C(=O)OCC(C)C |
| DIDP | CC(C)CCCCCCCOC(=O)C1=CC=CC=C1C(=O)OCCCCCCCC(C)C |
| DIHPP | CC(C)CCCCOC(=O)C1=CC=CC=C1C(=O)OCCCCC(C)C |
| DIHXP | CC(C)CCCOC(=O)C1=CC=CC=C1C(=O)OCCCC(C)C |
| DINP | CC(C)CCCCCCOC(=O)C1=CC=CC=C1C(=O)OCCCCCCC(C)C |
| DIOP | CC(C)CCCCCOC(=O)C1=CC=CC=C1C(=O)OCCCCCC(C)C |
| DITP | CC(C)CCCCCCCCCCOC(=O)C1=CC=CC=C1C(=O)OCCCCCCCCCCC(C)C |
| DIUP | CC(C)CCCCCCCCOC(=O)C1=CC=CC=C1C(=O)OCCCCCCCCC(C)C |
| DMP | COC(=O)C1=CC=CC=C1C(=O)OC |
| DNHP | CCCCCCOC(=O)C1=CC=CC=C1C(=O)OCCCCCC |
| DNPP | CCCCCOC(=O)C1=CC=CC=C1C(=O)OCCCCC |
| DPHP | CCCCCC(CCC)COC(=O)C1=CC=CC=C1C(=O)OCC(CCC)CCCCC |
| DPP | CCCOC(=O)C1=CC=CC=C1C(=O)OCCC |
| DNOP | CCCCCCCCOC(=O)C1=CC=CC=C1C(=O)OCCCCCCCC |
| DTDP | CCCCCCCCCCCCCOC(=O)C1=CC=CC=C1C(=O)OCCCCCCCCCCCCC |
| DUP | CCCCCCCCCCCOC(=O)C1=CC=CC=C1C(=O)OCCCCCCCCCCC |
| ODP | CCCCCCCCCCOC(=O)C1=CC=CC=C1C(=O)OCCCCCCCC |
| BPA | CC(C)(C1=CC=C(O)C=C1)C1=CC=C(O)C=C1 |
| BPADE | CC(C)(C1=CC=C(OCC2CO2)C=C1)C1=CC=C(OCC2CO2)C=C1 |
| BPAF | OC1=CC=C(C=C1)C(C1=CC=C(O)C=C1)(C(F)(F)F)C(F)(F)F |
| BPAP | CC(C1=CC=CC=C1)(C1=CC=C(O)C=C1)C1=CC=C(O)C=C1 |
| BPB | CCC(C)(C1=CC=C(O)C=C1)C1=CC=C(O)C=C1 |
| BPBP | OC1=CC=C(C=C1)C(C1=CC=CC=C1)(C1=CC=CC=C1)C1=CC=C(O)C=C1 |
| BPC | OC1=CC=C(C=C1)C(=C(Cl)Cl)C1=CC=C(O)C=C1 |
| BPE | CC(C1=CC=C(O)C=C1)C1=CC=C(O)C=C1 |
| BPF | OC1=CC=C(CC2=CC=C(O)C=C2)C=C1 |
| BPG | CC(C)C1=C(O)C=CC(=C1)C(C)(C)C1=CC(C(C)C)=C(O)C=C1 |
| BPM | CC(C)(C1=CC=C(O)C=C1)C1=CC(=CC=C1)C(C)(C)C1=CC=C(O)C=C1 |

| | |
|---|---|
| BPP | CC(C)(C1=CC=C(O)C=C1)C1=CC=C(C=C1)C(C)(C)C1=CC=C(O)C=C1 |
| BPPH | CC(C)(C1=CC(=C(O)C=C1)C1=CC=CC=C1)C1=CC=C(O)C(=C1)C1=CC=CC=C1 |
| BPS | OC1=CC=C(C=C1)S(=O)(=O)C1=CC=C(O)C=C1 |
| BPTMC | CC1CC(C)(C)CC(C1)(C1=CC=C(O)C=C1)C1=CC=C(O)C=C1 |
| BPZ | OC1=CC=C(C=C1)C1(CCCCC1)C1=CC=C(O)C=C1 |
| TBPA | CC(C)(C1=CC(Cl)=C(O)C(Cl)=C1)C1=CC(Cl)=C(O)C(Cl)=C1 |

[a] Abbreviations given in STable 1, *SMILES build by Marvin 17.21.0.*
